# Dual-mode ClfA-targeting DARPin biologics protect against diverse methicillin-resistant *Staphylococcus aureus* strains

**DOI:** 10.64898/2026.01.09.698026

**Authors:** Karuppiah Chockalingam, Biswarup Banerjee, Yu Zeng, Shuyu Miao, Joshua Bryant, Dominique Missiakas, Zhilei Chen

**Affiliations:** Department of Microbial Pathogenesis and Immunology, Texas A&M College of Medicine, Bryan, TX, USA; Department of Microbiology, Howard Taylor Ricketts Laboratory, The University of Chicago, Lemont, IL, USA; Genetics and Genomics Interdisciplinary Program, Texas A&M University, College Station, Texas, USA; Department of Biochemistry and Biophysics, Texas A&M University, College Station, Texas, USA

## Abstract

*Staphylococcus aureus* uses the adhesin clumping factor A (ClfA) to bind fibrinogen and promote invasive infection through two distinct interfaces: an exposed, low-affinity site on the N3 head domain and a buried, high-affinity “dock, lock, and latch” (DLL) trench that is exposed only under shear. This dual-interface architecture allows limited antibody penetration, as antibodies typically block only the exposed site. Here, we establish a dual-mode inhibition strategy that overcomes this constraint by combining a high-affinity ClfA-binding designed ankyrin repeat protein (DARPin) with a fibrinogen γ-chain peptide capable of occupying the DLL trench. Using cell-free click display and kinetics-guided affinity maturation, we engineer DARPin–γ-peptide fusion biologics that simultaneously block both fibrinogen-binding interfaces. These molecules inhibit ClfA–fibrinogen interactions, prevent methicillin-resistant *S. aureus* agglutination in human plasma, neutralize major clinical ClfA variants, and confer Fc-independent protection in a lethal murine bacteremia model. This work provides a strategy for targeting antibody-intractable force-activated staphylococcal adhesins.

## Introduction

*Staphylococcus aureus* (*S. aureus*) is a leading cause of bacteremia, endocarditis, osteomyelitis, and skin and soft-tissue infections^1^. Methicillin-resistant *S. aureus* (MRSA) is a dominant cause of healthcare-associated and, increasingly, community-acquired infections, ranging from skin abscesses to life-threatening bacteremia, endocarditis, osteomyelitis, and prosthetic-device infections^1, 2^. Current therapies for MRSA are facing increasing constraints including i) narrowing therapeutic window due to escalating resistance^3, 4^; ii) suboptimal drug penetration into abscesses, biofilms, and intracellular reservoirs^5, 6^; iii) prolonged IV administration for the most potent drugs^7, 8^; and iv) diminishing efficacy of existing drugs due to diverse resistance mechanisms^9, 10^. These concerns have prompted the CDC to classify MRSA as a high-priority public health threat and have underscored a critical need for non-antibiotic biologics that neutralize MRSA virulence factors essential for disease progression.

Clumping factor A (ClfA) is a promising *S. aureus* virulence factor for drug development. Present in >99% of clinical *S. aureus* isolates^11, 12^, ClfA is a surface-anchored microbial surface components recognizing adhesive matrix molecule (MSCRAMM) that binds host fibrinogen (Fg) and activated platelets, and drives the formation of fibrin-encased bacterial aggregates that protect MRSA from opsonophagocytosis and antibiotic penetration^13–16^. ClfA engagement of Fg is mediated by at least two distinct binding interfaces: a low-affinity site on the N3 head domain and a high-affinity trench between N2-N3 that recognizes the 17-residue C-terminal tail of the Fg γ-chain (γ-peptide) through an ultra-strong “dock, lock, and latch” (DLL) mechanism^17^. Notably, the DLL trench is buried under static conditions and becomes exposed only under shear stress^18^, enabling *S. aureus* to form mechano-responsive “catch bonds” and adhere avidly to Fg, Fg-coated surfaces and activated platelets in the vasculature^19–21^.

These structural and biophysical constraints have impeded the development of ClfA-targeted therapeutics. Monoclonal antibodies such as tefibazumab^14, 22^ and AZD7745 (formerly known as SAR114)^23, 24^ bind epitopes on the exposed N3 head and can only block the low-affinity Fg-binding site, but cannot access or competitively block stress-dependent interactions of Fg with the DLL trench. Moreover, antibodies raised against ClfA001 (ST8) often lack potency against ClfA002 (ST5), owing to genetic variations between strains (e.g. N463R) that disrupt antibody-binding without altering Fg-binding affinity^17, 25^. These limitations have contributed to the disappointing clinical performance of anti-ClfA antibodies^26, 27^, which have tended to be dependent on Fc-mediated opsonophagocytic killing and fail to fully suppress ClfA function^24–29^.

We hypothesized that simultaneous engagement of both the low- and high-affinity Fg-binding interfaces of ClfA by pairing a binder targeting the exposed N3 head with the Fg γ-peptide that can occupy the buried DLL trench should be a much more effective means of neutralizing *S. aureus*. Antibodies are ill-suited for this architecture. The Fg γ-peptide requires a free C-terminus and must be positioned proximally to the N3 epitope to optimally position it to thread the normally closed N2-N3 trench. This design is geometrically incompatible with large IgG scaffolds. In contrast, Designed Ankyrin Repeat Proteins (DARPins) are compact (12-18 kDa), modular molecules that can be easily engineered to position functional peptides adjacent to the binding interface, making them an ideal platform for dual-epitope blockage of ClfA.

In this work, we use a combination of cell-free high-diversity protein display (“Click Display”^30^) and selection for molecules that compete with Fg for ClfA-binding, followed by kinetics-focused affinity maturation and screening, to engineer ACE25-γ_D16A_, a DARPin-γ peptide fusion that engages both the N3 head and the DLL trench of ClfA. We further incorporate a human albumin-binding DARPin to generate two long-acting therapeutic candidates CADAR1 and CADAR2. These dual-action DARPin biologics are broadly effective against both ClfA001 (ST5) and ClfA002 (ST8) which collectively account for 60-100% of MRSA isolates^31^, showing both potent neutralization of the ClfA-Fg interaction and prevention of Fg-mediated MRSA agglutination in human plasma, regardless of the subtype. Finally, these molecules protect mice from lethal MRSA bacteremia, representing – to our knowledge – the first demonstration of Fc-independent protection against MRSA infection via direct ClfA neutralization.

Together, these results establish a new therapeutic design principle for MSCRAMM neutralization: functional mimicry of the host ligand (Fg γ-peptide) combined with high-affinity DARPin targeting of an adjacent structural domain. This modular strategy overcomes fundamental constraints inherent in antibody geometry, expands the reachable epitope space on force-activated bacterial adhesins, and provides a promising blueprint for next-generation anti-MRSA biologics.

## Results

### Discovery and engineering of 1^st^-generation ClfA-Fg inhibitory DARPins

Previously we developed a cell-free protein display technology – click display – that produces a covalently linked protein-cDNA complex from the template DNA in a one-pot reaction within 2 hours^30^ **(Figure 1A)**. A synthetic naïve DARPin library with up to ∼10^12^ variants (1 µg of template DNA) was produced using click display and an enriched pool of DARPins targeting ClfA001 N2-N3 (henceforth referred to as ClfA, **Figure S1**) was obtained after 6 rounds of selection^30^. In our previous study, all the cDNA bound to biotinylated ClfA (bClfA) and pulled down by streptavidin or neutravidin beads was amplified by PCR and used as template for the subsequent round of enrichment. Starting from the Round-6-enriched DARPin pool, in this study, we aimed to specifically select DARPins that bind at the interface of ClfA and Fg. Instead of amplifying all the cDNA pulled down by the beads, in Rounds 7 and 8 of panning, a high concentration of Fg (200 nM) was used to selectively elute DARPin-cDNA complexes from ClfA **(Figure 1B)**. We reasoned that DARPins that bind ClfA at a site which overlaps with that of Fg binding should be competitively eluted by Fg. The Round 7 DARPin pool showed slightly reduced background binding to streptavidin beads and slightly increased binding to ClfA, although the differences were not statistically significant **(Figure S2)**. The enriched DARPin pool from Round 8 were expressed in BL21(DE3) *E. coli* and the ability of 188 Round-8 DARPin clones in crude *E. coli* lysates to bind ClfA in the absence and presence of Fg (200 nM) was determined via ELISA **(Figure S3)**. Twenty-two DARPins showing an above-background ClfA-binding signal and >20% reduced binding signal in the presence of Fg were selected for further characterization.

**Figure 1.**
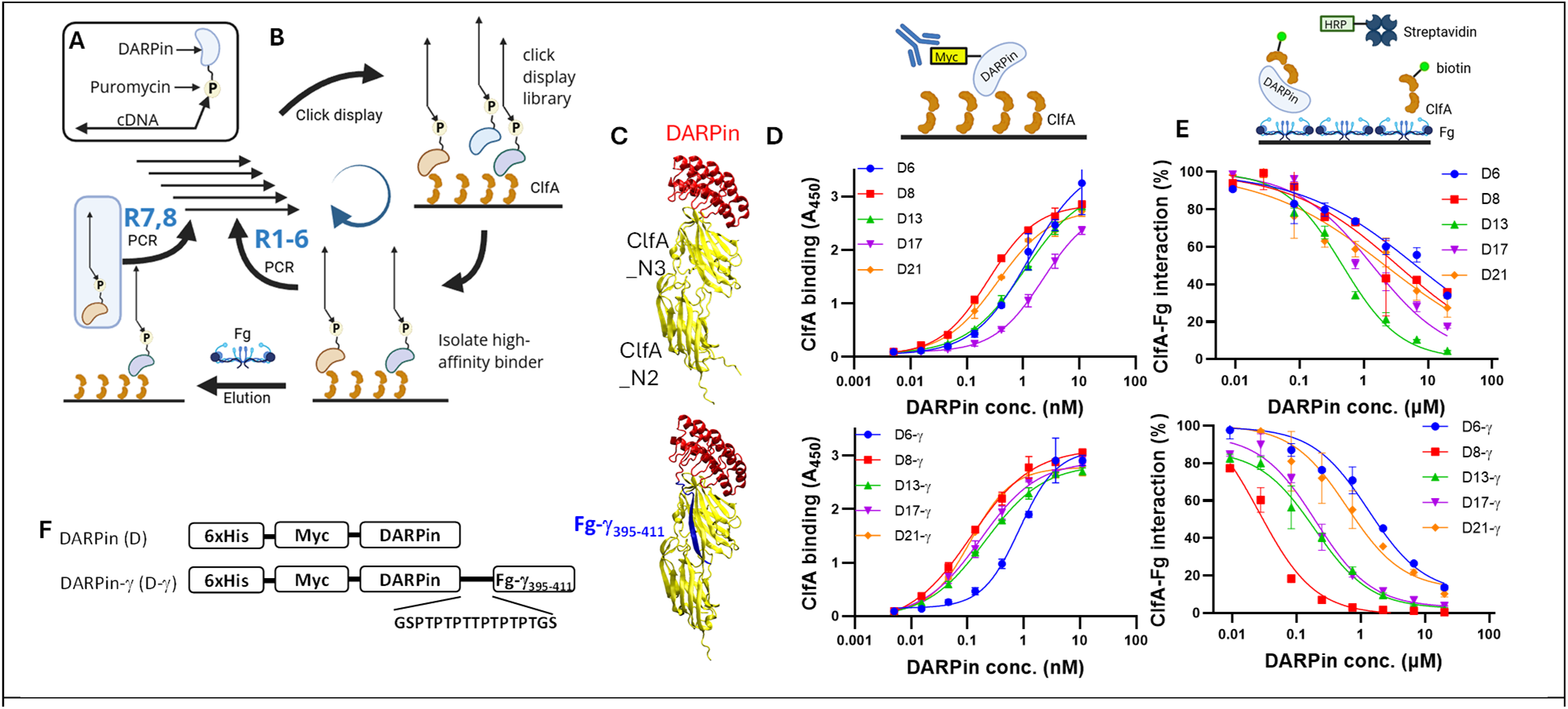
Characterization of 1^st^-generation anti-ClfA DARPins. **(A)** Schematic of click display. **(B)** Overview of click display library enrichment procedure. In selection Rounds 1-6, the cDNA of all ClfA-binding DARPins was amplified and used in the subsequent round. Starting from Round 7, only cDNA of DARPins competitively eluted by Fg was amplified. **(C)** AlphaFold2 model of D8 and D8-γ in complex with ClfA N2-N3. Yellow, ClfA; blue, Fg γ-peptide; red, DARPin D8. **(D)** ELISA binding titration of parent DARPins and DARPin-γ fusion constructs to immobilized ClfA. The ELISA plate was coated with streptavidin and bClfA, and DARPin binding was detected using anti-Myc antibody. **(E)** Dose-dependent inhibition of soluble ClfA binding to immobilized Fg by DARPin and DARPin-γ determined via competition ELISA. The ELISA plate was coated with Fg and incubated with 100 nM of bClfA and serially diluted DARPin or DARPin-γ molecules and plate-bound bClfA was detected using streptavidin-HRP. Data points are shown as mean ± SEM of duplicate readings from one of at least two independent, representative experiments. Curves were fitted using variable slope (three parameters) in GraphPad Prism. **(F)** Schematic of DARPin and DARPin-γ constructs.

Among these, five clones showed nanomolar affinity toward immobilized ClfA in ELISA **(Figure 1D, S4)**. The strongest ClfA binder, DARPin #8 (D8), exhibited a binding response EC_50_ of 0.3 nM **(Table 1)**. We next evaluated the ability of these DARPins to inhibit the interaction between soluble ClfA and immobilized Fg via competition ELISA. Briefly, Fg-coated ELISA wells were incubated with bClfA (final 100 nM) and serially diluted DARPin molecules (final 0-20 µM)^32^. After washing, the amount of bClfA in each well was quantified using streptavidin-HRP. All DARPins dose-dependently, albeit weakly, inhibited bClfA binding to Fg **(Figure 1E)**.

**Table 1.** Half maximal ClfA binding (EC_50_) and ClfA-Fg binding inhibitory (IC_50_) concentrations of 1^st^-generation anti-ClfA DARPins. Values were computed using GraphPad Prism (three parameters fit) from 2 independent experiments.

| | ClfA-binding $EC_{50}$ (nM) | | Fg-ClfA inhibitory $IC_{50}$ (μM) | |
| --- | --- | --- | --- | --- |
| | w/o $\gamma$ | w/ $\gamma$ | w/o $\gamma$ | w/ $\gamma$ |
| D6 | 1.5 ± 0.3 | 0.8 ± 0.003 | > 1 | 0.9 ± 0.6 |
| D8 | 0.3 ± 0.1 | 0.1 ± 0.01 | > 1 | 0.02 ± 0.005 |
| D13 | 1.1 ± 0.01 | 0.2 ± 0.03 | 0.5 ± 0.2 | 0.2 ± 0.02 |
| D17 | 2.4 ± 0.2 | 0.2 ± 0.002 | > 1 | 0.2 ± 0.07 |
| D21 | 0.4 ± 0.1 | 0.1 ± 0.02 | > 1 | 0.5 ± 0.1 |

All of these DARPins were predicted to bind to the head of the ClfA N3 domain by AlphaFold2 **(Figure 1C)**, with minor differences in the specific contact residues. We next fused the Fg γ-peptide (amino acids 395-411 of the Fg γ-chain^22^) to the C-terminus of each DARPin via a short PT-rich linker (∼30 Å)^33^ to generate a panel of DARPin-γ (D-γ) proteins **(Figure 1F, S4)**. All D-γ proteins showed enhanced ClfA binding ability compared to their respective parents **(Figure 1D**, **Table 1)** with D17-γ (EC_50_ 0.2 nM) exhibiting >10-fold lower EC_50_ relative to D17 (EC_50_ 2.4 nM). D8-γ showed the best Fg-inhibitory activity (IC_50_ 20 nM), at least one order of magnitude better than all other D-γ molecules, despite a modest (∼3-fold) enhancement in ClfA-binding affinity relative to the parent DARPin D8 (**Table 1**). The AlphaFold2 multimer model of D8-γ and ClfA showed Fg-like docking of the Fg γ-peptide into N2-N3 trench responsible for high-affinity Fg-binding (**Figure 1C**). We posit that different D-γ molecules may insert the Fg γ-peptide into the ClfA N2-N3 trench with varying efficiency depending on the precise ClfA-binding orientation of the DARPin.

### Affinity maturation of D8 to create 2^nd^-generation anti-ClfA DARPins

Encouraged by the superior Fg-inhibitory activity of D8-γ, we next set out to enhance the ClfA-binding affinity of the parent DARPin D8. The DARPin scaffold contains three internal ankyrin repeat domains (ARs) sandwiched between N- and C-Cap domains. In the original DARPin design, each AR contains six fully randomized target-binding residues – four in the loop and two on the α-helix – and one partially randomized residue located at the hinge position connecting to the loop. Our prior studies showed that residues surrounding the originally designated target binding residues can also play important roles in mediating target interaction^34^ and thus we expanded the number of fully randomized target-binding residues to eight in each AR and also randomized the loop region in the C-Cap in the naïve library (**Figure S4**). Error-prone PCR^35^ was used to introduce an average of 5-7 point mutations into the D8 gene and the resulting randomized library was subjected to three rounds of conventional click display selection followed by two rounds of off-rate selection^36, 37^ (**Table S1**). The three initial rounds of click-display selection were designed to select for DARPins with a higher kinetic on-rate and included a progressively increasing selection stringency in terms of (i) lower concentration of bClfA target protein, (ii) reduced incubation time of the click-displayed product library with the target prior to pull-down using streptavidin magnetic beads, and (iii) increased washing of the beads containing captured binders. For the off-rate selection, the click-displayed DARPin library was exposed to bClfA in the presence of a 500-1000-fold molar excess of the competitor molecule Fg to create a selective pressure for DARPins with a low off-rate.

The DARPin pool from the final round of off-rate selection was expressed in BL21(DE3) *E. coli* with an N-terminal Myc tag and C-terminal 6xHis tag. The ability of 147 individual clones in crude *E. coli* lysate to bind ClfA relative to the parent DARPin D8 was analyzed using biolayer interferometry (BLI) (**Figure S5**). Twenty-seven promising DARPin candidates from this preliminary screen were identified and purified by immobilized affinity chromatography (IMAC). Among these, eight unique DARPins showed >50-fold lower equilibrium dissociation constant (k_D_) compared to D8. Two of these clones – ACE6 and ACE25 –showed higher k_on_ and similar k_off_, while the remaining six showed higher k_on_ and lower k_off_, compared to D8 (**Figure 2A, S6**).

**Figure 2.**
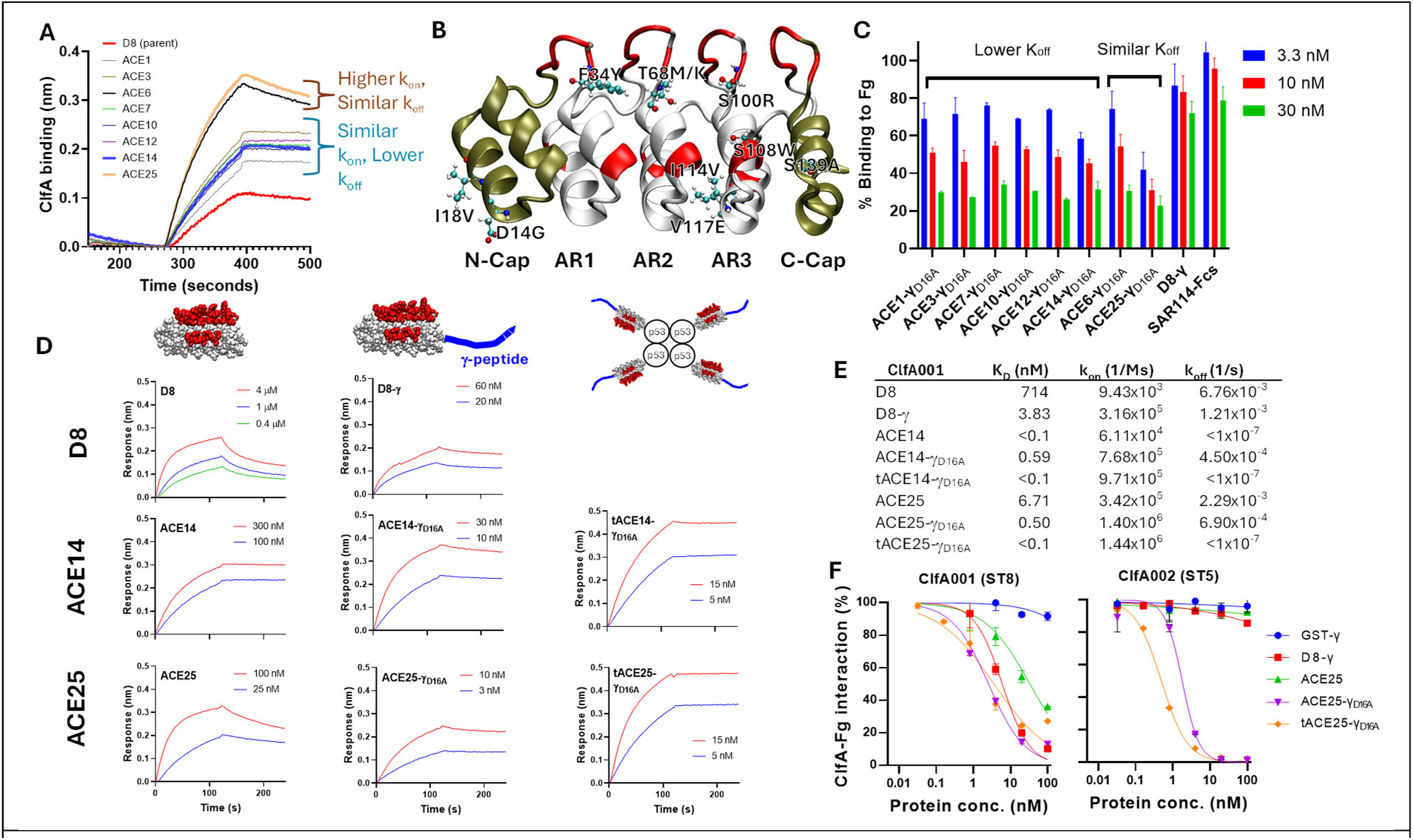
Evaluation of ClfA-binding and inhibition of Fg-ClfA binding by 2^nd^-generation anti-ClfA DARPins and their derivatives. **(A)** BLI sensograms of parent D8 and affinity-matured DARPin candidates. Streptavidin sensors were loaded with bClfA followed by 100 nM of each DARPin analyte. **(B)** Cartoon diagram of D8 predicted by AlphaFold2. Red: fully randomized residues in the naïve library; ball-and-sticks: sites and mutations present in affinity-enhanced anti-ClfA DARPins. **(C)** Dose-dependent inhibition of soluble ClfA binding to immobilized Fg by DARPin and DARPin-γ determined via competition ELISA. **(D)** ClfA-binding kinetics measured by BLI. **(E)** Summary of affinity data of different DARPins measured by BLI. **(F)** Dose-dependent inhibition of soluble ClfA binding to immobilized Fg by DARPin and DARPin-γ determined via competition ELISA. The ELISA plate was coated with Fg and incubated with 20 nM of bClfA001 (ST8) or 4 nM bClfA002 (ST5) and serially diluted DARPin or DARPin-γ. The binding of bClfA to immobilized Fg was detected using streptavidin-HRP.

Each of the eight leading affinity-matured DARPin candidates contains 3-6 amino acid substitutions relative to D8, many of which are at (F34Y, S100R, S108W) or immediately adjacent to (T68M/K, **Figure 2B**) DARPin sites designated for target binding. All of these DARPins contain the F34Y mutation, and all except ACE25 also contain the mutation S100R, indicating an important role of these substitutions in enhancing the ClfA-binding kinetics of D8 **(Figure S7)**. Since both positions 34 and 100 are fully randomized in the naïve DARPin library, these mutations likely directly contribute to improved ClfA-binding affinity. Six of the eight DARPins contain a substituted T68 residue (three to M and three to K), which is located immediately adjacent to the designated target-binding site 67 in AR2. DARPins ACE6 and ACE25 share identical mutations in AR1 and AR2. Our ability to identify multiple independent mutants with shared point-mutations reflects not only the importance of these positions/residues in enhancing ClfA-binding ability but also the power of the click display platform to efficiently produce large and diverse libraries that enable robust enrichment of target binders.

We next fused a D16A variant of the Fg γ-peptide, in which the penultimate aspartate residue is substituted with alanine, to the C-terminus of the 2^nd^-generation ClfA-binding DARPin candidates to form ACE_x_-γ_D16A_ analogous to D8-γ (**Figure 1F**). The D16A mutation has been reported to confer both increased binding affinity towards ClfA and decreased affinity towards the platelet integrin α_IIb_β_3_^22^, an interaction that under physiological conditions mediates Fg-induced platelet aggregation. The ability of ACE_x_-γ_D16A_ proteins to inhibit the interaction between soluble ClfA and immobilized Fg was evaluated using competition ELISA and compared to SAR114-Fcs, an Fc-silenced version of a broad-spectrum anti-ClfA antibody^25^ with picomolar ClfA-binding affinity, and the parent D8-γ protein (**Figure 2C**). All 8 ACE_x_-γ_D16A_ proteins showed stronger inhibition of the ClfA-Fg interaction than the parent D8-γ, which was comparable in inhibition potency to SAR114-Fcs. ACE25-γ_D16A_ and ACE14-γ_D16A_ showed the best Fg-inhibitory activity with >40% inhibition of ClfA binding at 3.3 nM. Next, we compared the binding affinity of different DARPins using biolayer interferometry (BLI). Binding kinetics measurements via a global fit of two or more biolayer BLI binding curves revealed a significant ClfA-binding enhancement conferred by fusion with the Fg γ-peptide (**Figure 2D,E**). ACE14 binds ClfA with >10^4^-fold lower k_off_ (<10^-7^ s^-1^) than D8 (6.8×10^-3^ s^-1^) while ACE25 exhibits a similar k_off_ (2.3×10^-3^ s^-1^) to D8 but significantly higher k_on_ (3.4×10^5^ Ms^-1^ for ACE25 vs. 9.4×10^3^ Ms^-1^ for D8). Interestingly, we observed different effects when fusing the Fg γ-peptide to these DARPins. Fusion of the wild-type Fg γ-peptide to D8 yielded >100-fold improvement in ClfA binding (K_D_ values of 714 nM for D8 and 3.8 nM for D8-γ). However, fusion of Fg-γ_D16A_ variant peptide to ACE25 enhanced ClfA binding only ∼13-fold (K_D_ values of 6.71 nM for ACE25 versus 0.5 nM for ACE25-γ_D16A_) while fusion of Fg-γ_D16A_ to ACE14 weakened the interaction by >5.9-fold (K_D_ values of 0.59 nM for ACE14-γ_D16A_ versus <0.1 nM for ACE14). We posit that the dampened enhancement conferred by fusion with the Fg γ_D16A_ peptide in the case of the ClfA-affinity-enhanced DARPins is suggestive of a “ceiling” being reached in ClfA binding affinity, beyond which additional affinity enhancement via synergy with the Fg γ-peptide DLL interaction is incremental or even impossible.

Since ClfA is a *S. aureus* bacterial cell wall-anchored protein, we reasoned that multimeric DARPins may exhibit better *in vivo* efficacy by simultaneously engaging multiple ClfA molecules in an aggregated/clumped complex. Tetrameric versions of the best ACE-γ_D16A_ proteins – ACE14-γ_D16A_ and ACE25-γ_D16A_ – were constructed by fusing a human p53 tetramerization domain (p53-TD) to the N-terminus to form tACE14 and tACE25-γ_D16A_. The p53-mediated homo-tetramerization conferred both tACE14-γ_D16A_ and tACE25-γ_D16A_ with significantly enhanced avidity to ClfA relative to the affinity of the respective parent monomeric protein, deriving primarily from a >4000-fold slower K_off_ rate.

During our entire DARPin engineering campaign, the N2-N3 domain of ClfA001 (ST8, Figure S1) was used as the target protein. In a new competition ELISA with immobilized Fg and lower concentrations of soluble ClfA001 (20 nM), ACE25-γ_D16A_ and tACE25-γ_D16A_ showed marginally better inhibitory activity than D8-γ **(Figure 2F)**. This may be due to the curves being “maxed out”, as 20 nM bClfA would theoretically require at least 20 nM DARPin to achieve 100% inhibition. Lower ClfA001 concentration yielded smaller signal-to-noise ratio and poor assay reproducibility. We also tested the inhibitory activity of these DARPins against the N2-N3 domain of ClfA002, a variant associated with highly virulent hospital-acquired sequence type 5 (ST5) strains^31, 38^ of *S. aureus* and significantly reduced affinity to anti-ClfA antibody tefibazumab^17^ and 11H10^25^. Both D8-γ and ACE25 showed marginal ClfA002-inhibitory activity at the highest tested concentration (100 nM). However, remarkably, both ACE25-γ_D16A_ and tACE25-γ_D16A_ showed strong ClfA002-inhibitory activity with estimated IC_50_ of 1.8 nM and 0.5 nM, respectively. Since neither ACE25 nor GST-γ-peptide significantly inhibited ClfA002 binding to immobilized Fg, the strong inhibitory activity of ACE25-γ_D16A_ and tACE25-γ_D16A_ is likely due to an avidity effect stemming from both the enhanced binding affinity of ACE25 as well as mutant Fg γ_D16A_-peptide. ST8 and ST5 strains together account for 60-100% of MRSA isolates from sepsis patients in the US between 2010 to 2022^31^. The ability of ACE25-γ_D16A_ and tACE25-γ_D16A_ to potently inhibit both ClfA001(ST8) and ClfA002(ST5) suggests that ACE25 likely binds a highly conserved epitope on ClfA and points to a high translational potential of these molecules.

### DARPins inhibit MRSA agglutination in human blood

Since purified N2-N3 domain of ClfA001 was used during DARPin engineering campaign, to ensure that these DARPins are also effective against full-length ClfA (N1-N2-N3) on the bacterial surface, we evaluated their ability to inhibit *S. aureus* agglutination. When inoculated in anti-coagulated human blood or plasma, the agglutination of *S. aureus* can be visualized as the assembly of large aggregates in a network of fibrin. *S. aureus* agglutination reaction requires bacterial factors ClfA as well as secreted Coa and vWbp, which collectively bind and activate prothrombin in the presence of fibrinogen^32, 39, 40^. This process closely represents the physiological host-pathogen interaction during *S. aureus*-induced sepsis^29^. Wild type Newman *S. aureus* (Newman) bacteria were fluorescently labeled with SYTO-9 and incubated in Na-Citrate human plasma in the presence or absence of different DARPins on glass slides for 30 minutes and imaged using a fluorescence microscope. The ClfA-binding antibody SAR114-Fcs and an isogenic *clfA* Newman mutant (*clfA*_Newman_) were included. Representative images are shown in **Figure 3A**. Newman formed large aggregates with average area of agglutinated bacteria of ∼5.5×10^7^ µm^2^, which is ∼100-fold larger than that *clfA*_Newman_ (7.6×10^5^ µm^2^, **Figure 3C**), highlighting a critical role of ClfA in *S. aureus* agglutination. All the DARPin constructs showed dose-dependent inhibition of *S. aureus* agglutination **(Figure 3A, B)**. The average areas of Newman aggregates when exposed to 5 nM and 50 nM of 2^nd^-generation DARPins (*i.e.* ACE25-γ_D16A_, tACE25-γ_D16A_, tACE14-γ_D16A_) are statistically indistinguishable from that of *clfA*_Newman_, indicative of complete neutralization of ClfA by these DARPin constructs. At 5 nM, the mean area of agglutinated Newman in the presence of the 1^st^-generation D8-γ was ∼10-fold larger than that for the 2^nd^-generation DARPins, regardless of p53-TD-mediated homo-tetramerization. tACE25-γ_D16A_ and ACE25-γ_D16A_ were chosen for further characterization.

**Figure 3.**
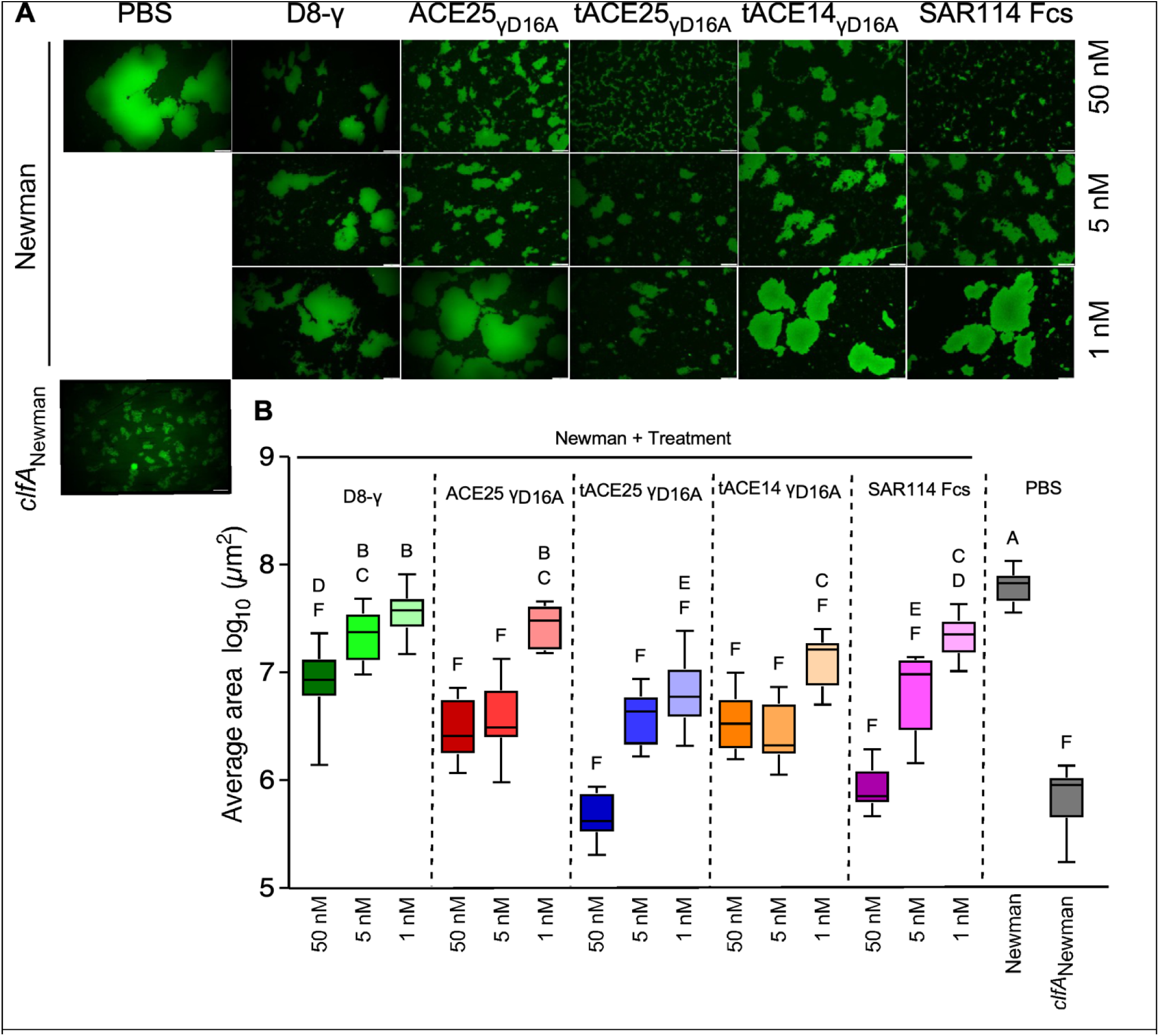
DARPins inhibit *S. aureus* agglutination in anti-coagulated human plasma. **(A)** Representative images of SYTO-9-stained Newman bacteria agglutinated in human plasma with either anti-ClfA DARPins or SAR114-Fc (positive treatment) and PBS (negative treatment). *clfA*_Newman_ was used as benchmark for reduced agglutination. Images were observed under an inverted fluorescence microscope with a 20× objective. **(B)** Box and whisker plot representation of staphylococcal agglutination areas with anti-coagulated human plasma observed in 10 different fields of microscopic view. Statistical significance was calculated in pairwise comparison using Brown-Forsythe and Welch ANOVA followed by Dunnett s T3 correction for multiple comparisons. Letters denote statistical significance as observed by multiple comparisons across samples. Experiment was performed twice for reproducibility.

The Newman strain belongs to the sequence type 8 (ST8) and harbors a ClfA001-type sequence. To gain insights into the broadness of our DARPins the agglutination assay was repeated with another ST8 strain USA300 LAC (herein referred to as USA300) and its isogenic *clfA* mutant (as a negative control, *clfA*_USA300_) and the ST5 isolate, N315. Similarly potent agglutination inhibition was observed for tACE25-γ_D16A_ and ACE25-γ_D16A_ against both wild type strains; USA300 and N315 bacteria treated with 50 nM of either DARPin formed very small clumps indistinguishable from the negative control *clfA* _USA300_ mutant strain **(Figure S8)**.

### DARPins protect mice from MRSA

Due to their small size, DARPins are mostly excreted within 7.5 minutes in mice upon *i.v.* injection^41^. Several strategies have been developed to extend the *in vivo* half-life of DARPins, including PEGylation^41, 42^, piggybacking on serum albumin^43, 44^ and fusion to unstructured polypeptides^45, 46^. Previously, the fusion of two copies of a serum albumin-binding DARPin (DA50) was found to significantly improve the circulation half-life of DARPins to ∼2 weeks in humans^47, 48^, prompting us to use the same serum-albumin-binding DARPin-assisted half-life extension strategy in constructs α**C**lf**ADAR**Pin**1** (CADAR1) and CADAR2. CADAR1 is a homo-tetramer with DA50 at the N-terminus of tACE25-γ_D16A_ while CADAR2 is a linear molecule with two tandem copies of DA50 inserted at the N-terminus of ACE25-γ_D16A_ **(Figure 4A)**. The fusion of the serum albumin-binding DARPin DA50 conferred both CADAR1 and CADAR2 with a relatively long half-life, with the linear CADAR2 showing both a higher maximum plasma concentration and longer systemic half-life compared to the homo-tetrameric CADAR1 (**Figure 4B**). Both proteins were detectable in the mice plasma 7 days following intraperitoneal injection.

**Figure 4.**
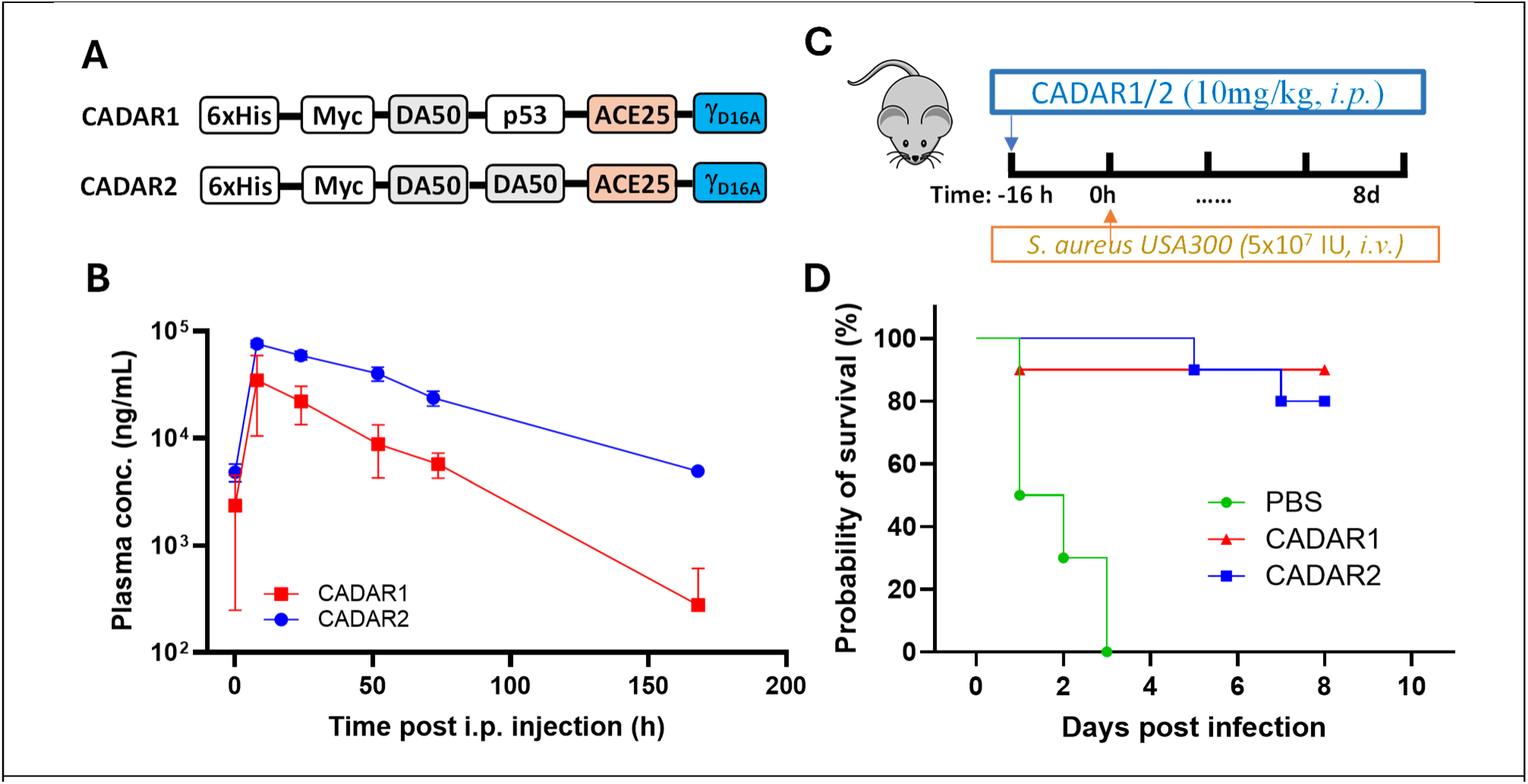
*In vivo* characterization of CADAR1 and CADAR2. **(A)** Schematic of CADAR1 and CADAR2. **(B)** Plasma concentration time course in C57BL/6 mice following intraperitoneal (i.p.) injection at a 10 mg/kg dose. Shown is the mean +/– SD of n=3 (CADAR1) or n=2 (CADAR2) experimentally independent animals for each time-point and treatment. **(C)** Schematic of *S. aureus* lethal intravenous (i.v.) challenge experiment. **(D)** Survival of mice challenged with a lethal dose (5 × 10^6^ CFU) of *S. aureus* USA300 after treatment with CADAR1 or CADAR2.

Next, we evaluated the efficacy of CADAR1/2 using a lethal *S. aureus* challenge model. Mice were passively immunized with 10 mg/kg CADAR1/2 via *i.p.* injection and, 16 hours later, retro-orbitally challenged with a lethal dose (5 × 10^6^ Colony Forming Units, CFU) of *S. aureus* USA300. All mice receiving PBS died within 3 days, while 9/10 and 8/10 mice receiving CADAR1 and CADAR2, respectively, remained alive at the end of the study (8 days post infection). In previous studies utilizing candidate anti-ClfA antibody therapeutics, opsonophagocytic killing (OPK) of bacteria mediated by the Fc domain was found to be critical for *in vivo* efficacy^24^. Since our DARPins lack an Fc component, the *in vivo* efficacy of CADAR1 and CADAR2 derives solely from the potent Fg-inhibitory activity, likely enhanced by the simultaneous blockade of both low- and high-affinity Fg-binding interfaces on ClfA.

## Discussion

Clumping factor A (ClfA) is a critical mediator of *S. aureus* virulence and a compelling therapeutic target due to its essential roles in fibrinogen binding, platelet activation, immune evasion and clot-embedded bacterial survival. Although its importance has been recognized for more than two decades, ClfA has remained challenging to neutralize effectively *in vivo*. This difficulty stems from its dual-mode fibrinogen (Fg) engagement – a low-affinity binding surface on the N3 head domain and a high-affinity, shear-activated “dock, lock, and latch” (DLL) trench between N2 and N3. Since natural anti-ClfA antibodies can only target one surface, and the DLL trench is hidden under static conditions, existing evidence suggests that antibodies most often neutralize the N3 domain^22, 23^, leaving the most biologically potent fibrinogen-binding mechanism intact. Moreover, several ClfA variants – including ClfA002 – escape antibody recognition despite retaining Fg binding, further complicating antibody-based approaches^17, 24, 25^.

Our work addresses these long-standing limitations by establishing a dual-mode DARPin-γ-peptide fusion architecture capable of simultaneously blocking both Fg-binding interfaces on ClfA. Through click display and functional screening for molecules that bind an overlapping epitope with Fg, we identified a panel of DARPins that dose-dependently inhibit the ClfA-Fg interaction. Fusion of the Fg γ-peptide to one of these DARPins – D8, predicted to bind the N3 head domain – yielded D8-γ which facilitated the positioning of a free C-terminal Fg γ-peptide within the DLL trench. This dual-site engagement conferred >50-fold enhanced inhibition of the ClfA-Fg interaction compared to the parent D8 **(Table 1)**, demonstrating an effective structure-guided strategy for accessing epitopes that are not easily accessed or transiently exposed under physiological shear.

An important advantage of our strategy is its tunability and modularity. Unlike antibodies, whose geometric constraints prevent the placement of a free C-terminal γ-peptide in close proximity to the N3 epitope, DARPins provides a compact scaffold enabling flexible spatial alignment of the peptide ligand. This architecture was further strengthened by affinity maturation to adjust kinetic parameters – particularly the on-rate – to promote rapid docking onto ClfA.

The parent D8 and D8-γ molecules showed negligible Fg-inhibitory activity toward ClfA002 (ST5), a result which is not surprising given that the N2-N3 domain of ClfA001 (ST8) was used as the panning target during the entire DARPin engineering campaign and the ClfA002 strain harbors a N463R mutation at the N3 head domain relative to ClfA001. Similar results have been observed in the antibody field, with ClfA001 N2-N3 immunization yielding antibodies – tefibazumab and 11H10 – showing significantly reduced affinity to ClfA002. Tefibazumab (derived from mAb 12-9^26^) was found to bind a ClfA_N463R_ isoform with a 60-fold reduced affinity^17^ while 11H10 showed >40-fold reduced Fg-inhibitory potency toward ClfA002 relative to ClfA001^24, 25^. Upon affinity maturation and fusion to a D16A-mutant Fg-γ peptide, the resulting ACE25-γ_D16A_ molecule exhibits strong Fg-inhibitory activity toward both ClfA001 and ClfA002. In a MRSA agglutination assay using human plasma, a physiologically relevant model that integrates ClfA, secreted coagulases (Coa, vWbp), Fg and fibrin polymerization, ACE25-γ_D16A_ and its tetrameric derivative effectively reduced aggregate sizes to those of a *clfA* mutant, demonstrating near-complete neutralization of ClfA-mediated adhesion of intact bacteria at 50 nM concentration **(Figure 3)**. Critically, this effect extends to multiple MRSA strains, including USA300 and N315 **(Figure S8)**, indicating broad translational potential.

By incorporating a serum albumin-binding DARPin (DA50)^47, 48^, we generated CADAR1 and CADAR2, long-lived therapeutic candidates with distinct architectures but comparable activity. In a lethal USA300 bacteremia model, a single dose of either molecule conferred 80-90% survival, despite the absence of an Fc domain **(Figure 4)**. This finding is particularly notable as prior work has shown that anti-ClfA antibodies are dependent on Fc-mediated opsonophagocytosis for effective *in vivo* protection, and antibodies lacking complement and Fcg receptor engagement lose significant activity even when they strongly inhibit the ClfA-Fg interaction^24^. In contrast, CADAR1/2 confer strictly Fc-independent protection, demonstrating that direct neutralization of ClfA s adhesive function is sufficient to significantly reduce bloodstream lethality. This Fc independence may be especially advantageous in the context of *S. aureus*, which has evolved multiple mechanisms – such as Protein A – to block antibody effector function.

Our data also reveal potential avenues for further optimization. Although tetramerization enhanced avidity, the short linker connecting DARPins within CADAR1 and tACE25-γ_D16A_ may limit simultaneous engagement of multiple spatially separated ClfA molecules on the bacterial surface. Systematic optimization of linker length, rigidity and/or geometry may further improve avidity-driven neutralization. Moreover, ClfA interacts not only with Fg but also with complement regulator Factor I^49–51^. ACE25-γ_D16A_ does not inhibit this interaction **(Figure S10)**, suggesting that the Factor I and Fg binding sites are distinct. Additional engineering efforts geared towards simultaneous blockade of Fg and Factor I to ClfA may yield therapeutically more efficacious molecules.

More broadly, this work demonstrates a generalizable principle: DARPin-tethered ligand mimicry provides a powerful means to target geometrically occluded or force-activated epitopes that may be inaccessible to antibodies. This approach may be extended to other MSCRAMMs and adhesins that undergo conformational changes under physiological shear. Because DARPins are compact, modular, highly stable, and have demonstrated generally low immunogenicity in clinical settings to date, long-acting DARPin-γ fusion biologics may be well-suited for prophylaxis or adjunctive therapy in high-risk populations.

In summary, we have introduced a new class of dual-mode ClfA inhibitors that directly neutralize the two Fg-binding interfaces on ClfA essential for MRSA pathogenesis. By overcoming intrinsic limitations of antibody geometry and targeting adhesin mechanics that are central to *S. aureus* bloodstream infection, these DARPin-templated molecules represent a promising new modality for anti-MRSA biologics. Future work will explore therapeutic dosing, efficacy in additional infection models, epitope mapping, and structural characterization to further refine this platform and expand its application to other virulence factors.

## Methods

### Plasmid construction

All DARPin constructs were sub-cloned into pET28a downstream of a T7 promoter and contain an N-terminal c-Myc tag and a 6xHis tag at N-/C-terminus. For DARPin-γ fusion constructs, a gene fragment encoding the Fg-γ_395-411_ peptide and upstream linker was synthesized by Integrated DNA Technologies and appended to the C terminus of the selected DARPins. For constructs containing the D16A variant of the Fg γ-peptide, the penultimate aspartate residue was substituted with alanine by overlap extension PCR^52^.

To construct tetrameric forms of DARPin-γ, a gene fragment (GenScript) encoding the human p53 tetramerization domain (p53-TD) followed by a (GGGGS)_2_ linker was inserted upstream of the DARPins. Protein sequences for CADAR1 and CADAR2 are provided in **Figure SG**.

### Preparation of non-DARPin proteins reagents (ClfA, fibrinogen, antibody, filler protein)

The N2-N3 portion of ClfA-001 (ST8, residues 221 – 561; referred to as “ClfA” in this study unless otherwise indicated) and ClfA-002 (ST5, residues 221-561) was expressed and purified by the Protein Production, Characterization, and Molecular Interaction (PPCMI) core facility at the Institute of Biosciences and Technology (IBT), Texas ACM University. Full sequences are provided in **Figure S1**. For biotinylation of ClfA, 100 μM ClfA in PBS was incubated with 400 μM freshly prepared EZ-Link Sulfo-NHS-LC-Biotin (Thermo Fisher) at room temperature for 30 minutes followed by 4°C overnight. Unreacted biotin was removed using two consecutive 0.5-mL 7K MWCO Zeba Desalting Columns (Thermo Fisher). Human fibrinogen (Fg) was a kind gift of Prof. Magnus Hook^53^. SAR114-Fcs (**Figure S1**), a variant of the anti-ClfA monoclonal antibody SAR114^25^ in which the Fc effector functions were silenced by a combination of LALAPG^54^ and the glycosylation-preventing N297Q mutation^55^ was expressed in Expi293F cells (Thermo Fisher) and purified using Protein A resin as described^56^. Biotinylated filler protein for blocking unoccupied sites in streptavidin beads was prepared by incubation of 133 μM hAC11-10^57^ (IgG1) in PBS with 800 μM freshly prepared EZ-Link Sulfo-NHS-LC-Biotin at room temperature for 30 minutes then at 4°C for 3 h, followed by cleanup and exchange into PBS using two consecutive 7K MWCO Zeba Desalting Columns.

### Identification of 1^st^-generation ClfA-Fg inhibitory DARPins via click display

Previously we carried out 6 rounds of click-display selection for DARPins that bind ClfA^30^. In this study (Rounds 7, 8), the output of Round 6 selection campaign was PCR-amplified and 1 µg of DNA was used as template for click display as described previously^30^. The DARPin molecule contains a 6xHis tag at the N-terminus. The click-displayed DARPin-cDNA was purified by one-step IMAC and incubated with bClfA (final 200 nM) in SBTD0.3 buffer (StartingBlock™ Blocking Buffer (Thermo Fisher) supplemented with 0.05% Tween 20 and 0.3 mg/mL ssDNA (Deoxyribonucleic acid sodium salt from salmon testes, Sigma) for 1 hour. Next, DynaBeads MyOne Streptavidin T1 magnetic beads (Thermo Fisher; streptavidin beads) were added to pull-down bClfA and associated DARPin-cDNA. After thorough washing, the beads were incubated in 1X Q5 reaction buffer (NEB) supplemented with 200 nM Fg for 30 minutes. The supernatant was collected. The beads were washed once with 1X Q5 reaction buffer and combined with the supernatant. The mixture was concentrated by ethanol precipitation and PCR-amplified using Q5 DNA polymerase (NEB) and primers 1938/2643. The PCR product was gel-purified and used as the template for the subsequent round of selection.

The PCR product encoding enriched DARPin pool from Round 8 was sub-cloned into the pET28a expression vector and used to transform BL21(DE3) cells. All DARPins harbor an N-terminal 6xHis tag and Myc-tag. One hundred and eighty-eight clones were picked and their ability to bind ClfA in the absence and presence of Fg (200 nM) was determined via ELISA **(Figure S3)**. Briefly, individual *E. coli* colonies were grown in 200 µL LB in flat-bottom 96-well plates at 37°C with shaking (400 rpm) overnight. The next day, 10 µL of the overnight culture from each well was transferred to fresh wells containing 190 µL of growth medium and grown at 37°C with shaking (525 rpm). The cultures were induced at OD_600_ 0.5, with 1 mM IPTG at 37°C for another 4 hours prior to centrifugation and storage of pellets at - 80°C. To prepare lysates, each pellet was resuspended in 100 µL lysis buffer (PBS with 0.2 mg/mL lysozyme, 1 mM CaCl_2_, 0.5 mM EDTA and 25 U/mL benzonase (Millipore), incubated at 37°C for 30 minutes, followed by three freeze-thaw cycles between -80°C and 37°C.

For functional screening, the ELISA plates were coated with 4 µg/mL ClfA001 in PBS overnight at 4°C. The next day, the wells were blocked with PBSTB (PBS w/ 0.1% Tween-20 and 2% BSA) and washed with PBST (PBS w/ 0.1% Tween-20). Next, DARPin-containing lysates were diluted 3.5-fold with PBS in the presence of 200 nM Fg and 140 μL of these mixtures were added to the wells. The ELISA plates were incubated at room temperature for 1 hour, followed by washing and detection of plate-bound DARPin using mouse anti-Myc primary antibody (Invitrogen), goat anti-mouse horseradish peroxidase (HRP) secondary antibody and 3,3,5,5 -tetramethylbenzidine (TMB; Surmodics) substrate **(Figure S3)**.

### Affinity maturation of DARPin D8

Error-prone PCR^35^ was used to create a point mutagenized library templated on DARPin D8 and this library (epDARPin-cDNA) was initially subjected to 3 rounds of conventional click-display selection essentially as described previously^30^ using a progressively decreasing concentration bClfA as the target protein for panning (**Table S1**). The crude click display products were used directly for selection and all dilution, blocking, and wash steps utilized SBTD0.6 (StartingBlock™ Blocking Buffer supplemented with 0.05% Tween 20 and 0.6 mg/mL ssDNA (Deoxyribonucleic acid sodium salt from salmon testes). Round 1 of selection utilized 20 nM bClfA as the target and the beads containing magnetically captured library molecules were washed 6 times. Round 2 used 10 nM bClfA with 8 washes and Round 3 employed 2.5 nM bClfA with 12 washes. In Round 3, in order to further minimize non-specific binding of library to the streptavidin beads, the beads containing the captured library was incubated with 500 nM of an irrelevant biotinylated “filler protein”, an IgG, at room temperature for 30 minutes before the 7^th^ wash. The filler protein is intended to displace library molecules that adhere non-specifically to unoccupied streptavidin sites on the beads through weak interactions. We next carried out 2 rounds of off-rate selection. Briefly, the click-displayed DARPin library was first incubated with naked streptavidin beads in SBTD0.6 at room temperature for 1 h to remove library molecules that bind non-specifically to the beads. The supernatant was then incubated with 1 nM bClfA at room temperature for 5 minutes (Round 4) or 1 minute (Round 5) prior to the addition of 500 nM Fg. The mixture was incubated at RT with rotation for 1 h (Round 4) or overnight (Round 5). In Round 5, after overnight incubation, the off-rate selection mixture was supplemented with an additional 500 nM Fg (1000 nM final Fg concentration) and incubated at room temperature for 4 hours followed by 37°C for 2 hours. Following the incubation with Fg, streptavidin beads were added to pull-down bClfA and associated epDARPin-cDNA. The beads were washed 12 times, with the 7^th^ wash including incubation with biotinylated filler protein at RT for 30 minutes as described above.

The final enriched DARPin pool was amplified from the streptavidin beads and sub-cloned into the pET28a expression vector in-frame with an N-terminal Myc tag and C-terminal 6xHis tag, expressed in BL21(DE3) cells, and grown in deep-96-well plates for BLI-screening of lysates as described in **Figure S5**.

### Biolayer interferometry (BLI)

Screening of DARPin-containing *E. coli* lysates was performed on an Octet Red96e instrument (ForteBio/Sartorius) using streptavidin (SA) sensors. All kinetics measurements were performed on a ForteBio BLItz (Octet N1, Sartorius) instrument using SA sensors. All proteins were diluted in Assay Buffer (PBS, 0.02% Tween-20, 0.1% BSA). DARPin-containing *E. coli* lysates were diluted 1000-fold in Assay Buffer. After sensor hydration in Assay Buffer for 10 mins, the sensor was loaded with 100 nM (lysate screening) or 50 nM bClfA (kinetics measurements). A single set of 8 sensors was loaded only once with bClfA and re-used for the entire lysate screening campaign. Re-use of sensors was enabled by low-pH regeneration with 10 mM Glycine-HCl, pH 2.0 between successive exposures to new lysate samples as described in Figure S5. Fresh SA sensors were used for each kinetics curve measurement.

### Protein expression and purification

For initial characterization of individual DARPin and DARPin-γ clones derived from click-display selection, the DARPin constructs in BL21 *E. coli* were grown in 5-mL Luria-Bertani (LB) Broth at 37°C OD_600_ until mid-log phase (OD_600_ 0.5 – 0.8) and induced with IPTG at 37°C for 4 hours prior to cells harvest. Cell pellets were lysed using B-PER II Bacterial Protein Extraction Reagent (Thermo Fisher) and the proteins purified by IMAC using 30-μL of settled Ni-NTA agarose (Cube Biotech) with Pierce 0.8-mL centrifuge columns (Thermo Fisher). DARPins were eluted twice using 20 mM sodium phosphate, pH 7.1, containing 250 mM imidazole, and the pooled eluate was immediately exchanged into PBS using 0.5-mL 7K MWCO Zeba desalting columns. Purified proteins were stored at 4°C for up to a week or supplemented with 50% glycerol at -20°C. One-liter BL21 cultures containing CADAR1 and CADAR2 were similarly induced with IPTG followed by lysis of cell pellets via sonication (QSonica Q500) and IMAC purification using 4-mL settled Ni-NTA agarose with 20-mL gravity-flow columns (G-Biosciences). Purified proteins were concentrated and exchanged into PBS using 50-kDa (CADAR1) or 30-kDa (CADAR2) Amicon Ultra-15 ultracentrifugal devices (Merck Millipore). The final purified CADAR1 and CADAR2 were treated to remove endotoxins using 1-mL Pierce High-Capacity Endotoxin Removal Columns (Thermo Fisher), sterilized using 0.2 μm PES filtration, aliquoted, and flash-frozen prior to storage at -80°C.

### Creation of DARPin-γ fusion constructs

A gene fragment encoding the Fg-γ_395-411_ peptide and upstream PT-rich linker was synthesized by Integrated DNA Technologies and appended to the C terminus of selected DARPins via a BsaI (Type IIs) restriction site introduced at the junction. The fusion construct including in-frame 6xHis and c-myc tags at the N terminus was sub-cloned with flanking BamHI and HindIII restriction sites into the pET28a expression vector downstream of a T7 promoter. For constructs containing the D16A variant of the Fg γ-peptide, the penultimate aspartate residue was substituted with alanine by overlap extension PCR^52^.

### ClfA-binding and ClfA-Fg inhibition ELISA

To evaluate the ability of DARPins to bind ClfA, ELISA MAX 96-well plates (BioLegend) were coated with streptavidin (4 µg/mL, Thermo Fisher) in phosphate-buffered saline (PBS) overnight at 4°C, blocked with PBSTB (PBS, 0.1% Tween-20, 2% bovine serum albumin) at room temperature (RT) for 1 hour and then incubated with bClfA (ST8 or ST5). After three washes (PBS, 0.1% Tween-20), DARPins diluted in PBSTB were added. The plates were incubated at room temperature for 30 minutes. Following three washes, bound DARPin was detected using HRP-conjugated chicken anti-Myc antibody (Immunology Consultants Laboratory) and TMB substrate. The absorbance at 450 nm (A_450_) was read on a Tecan Infinite M200 plate reader.

To determine the ability of DARPins to inhibit the Fg-ClfA interaction, ELISA plates were coated with 2 μg/mL Fg in PBS at 4°C overnight with rocking, washed 3 times, and blocked for 1 h at RT with PBSTB or StartingBlock buffer (Thermo Fisher) containing 0.1% Tween-20 (SBT; for CADAR1 and CADAR2 only). The plate was incubated with a mixture of 100 nM bClfA and serially diluted anti-ClfA proteins in a 100-μL final volume of PBSTB or SBT (CADAR1, CADAR2 only) for 1 h at RT. After 3 washes, bound bClfA was detected using HRP-streptavidin (1:20,000; Thermo Fisher) and 100 μL TMB. The reaction was stopped after approximately 6 mins with 100 μL 1 M H_2_SO_4_. The absorbance at 450 nm (A_450_) was read on a Tecan Infinite M200 plate reader.

The percentage inhibition of the ClfA-Fg interaction was determined using the following formula: 100 – [100 x ((A_450, ClfA+DARPin_ – A_450, No protein_)/(A_450, ClfA, No DARPin_ – A_450, No protein_))].

### Pharmacokinetics of CADAR1 and CADAR2

Mouse studies were performed according to Texas ACM University IACUC guidelines (protocol # 2023-0225). Eight-week-old BL57/6 mice were injected i.p. with 10 mg/kg of CADAR1 or CADAR2 in PBS at 10 mg/kg dose in 100 μL volume. Blood was collected by tail nick 5 minutes, 8 h, and 1, 2, 3, and 7 days following injection and immediately diluted 3-fold with PBS/10mM EDTA. Samples were further diluted with StartingBlock (Thermo Fisher) then centrifuged at 1500 x g for 10 mins to pellet red blood cells. The diluted plasma was flash frozen and stored at -80°C. The plasma concentration of CADAR1/2 was determined using ClfA-binding ELISA as described above, except that SBT was used instead of PBSTB for plate blocking and dilution steps.

### S. aureus agglutination assay

Staphylococcal agglutination in human plasma was measured as described^39^. 10 ml overnight cultures of *S. aureus* USA300 and *clfA* isogenic mutant, or N315 wild type were centrifuged, washed with PBS and resuspended in PBS at OD600 nm=4. SYTO 9 (Invitrogen) was added to the bacterial suspension for 15 mins in dark and washed twice with PBS to remove excess stain and the pellet was resuspended in 1 ml PBS. 1, 5 or 50 nM DARPins were added to the bacterial suspensions and PBS treatment was kept as control wherever indicated. Then equal volumes of pre-treated bacterial suspensions and anti-coagulated human plasma were added on to glass slides followed by 30 min incubation at room temperature. Bacterial agglutinates were observed under an inverted fluorescence microscope (Olympus). Agglutinated areas were measured from 10 different fields of view using QuPath software. Data were represented as box and whisker plot with standard error of mean.

### S. aureus lethal challenge

6-8 weeks old BALB/cJ mice were pre-treated with either PBS or 10 mg/kg CADAR1 and CADAR2 intraperitoneally. After 16 hours, mice were anesthetized with isoflurane, and a lethal dose of 5 × 10^6^ CFU of *S. aureus* USA300 was injected intravenously via retro-orbital route. Survival of the mice were observed for 8 days and data were plotted as probability of survival (Kaplan-Meier curve). The experiment was performed twice for reproducibility. These studies were performed according to the University of Chicago IACUC guidelines.

## Abbreviations

ClfA: clumping factor A;
bClfA: biotinylated ClfA
MRSA: Methicillin-resistant *S. aureus*
DLL: dock, lock and latch
Fg: fibrinogen
MSCRAMM: microbial surface components recognizing adhesive matrix molecule
BLI: biolayer interferometry
p53-TD: p53 tetramerization domain;

## Acknowledgements

We thank Ivantha Nawaratna, Protein Chemistry Lab, College of Agriculture and Life Sciences, Texas ACM University and Rodrigo Jacamo, Sartorius Corporation, for hands-on help with settings and use of the Octet Red96e instrument, and Wen Liu, Protein Production, Characterization, and Molecular Interaction Core at Texas ACM Health Institute of Biosciences and Technology, for providing purified recombinant ClfA proteins. We also thank Prof. Magnus Hook who inspired and encouraged us to engineer DARPin biologics against ClfA. This work was supported by Texas ACM Health Science Center Seedling grant, Texas ACM ADM grant and NIH R21AI182950.

## Author contributions

Z.C. and K.C. conceptualized the project. K.C. and Z.C. wrote the first draft of the manuscript with input from all authors. Z.C, K.C., D.M. and B.B. designed the experiments, led the analysis and contributed to writing the manuscript. Z.C. and D.M. supervised the research. K.C., Y.Z., B.B., S.M. and J.B. performed wet laboratory experiments. Z.C. designed and Y.Z. designed, performed and analyzed the experiments for the 1^st^-generation DARPin discovery campaign. K.C. and Z.C. designed, and K.C. performed the 2^nd^-generation DARPin discovery campaign, including affinity maturation and construction/evaluation of tetrameric and half-life-extended DARPin derivatives. D.M. designed and B.B. designed, performed and analyzed the human plasma agglutination assays and mouse studies with MRSA. S.M. performed ELISA characterization assays of 2^nd^-generation anti-ClfA DARPins. J.B. performed DARPin pharmacokinetics studies.

## Data availability

All oligonucleotide and protein sequences are included in the Supplementary Information. Statistics and plots were generated using GraphPad Prism v.8 (GraphPad Software, LLC). Additional source data are available from the corresponding authors upon reasonable request.

