## Supplementary Information for "Dual-mode ClfA-targeting DARPin biologics protect against diverse methicillin-resistant *Staphylococcus aureus* strains"

*ClfA-001 (ST8) N2-N3 sequence (after TEV protease removal of N-terminal 6xHis tag)*

**GS**VAADAPAAGTDITNQLTNVTVGIDSGTTVYPHQAGYVKLNYGFSVPNSAVKGDTFKITVPKELNL
NGVTSTAKVPPIMAGDQVLANGVIDSDGNVIYTFTDYVNTKDDVKATLTMPAYIDPENVKKTGNVTL
ATGIGSTTANKTVLVDYEKYGKFYNLSIKGTIDQIDKTNNTYRQTIYVNPSGDNVIAPVLTGNLKPNTD
SNALIDQQNTSIKVYKVDNAADLSESYFVNPENFEDVTNSVNITFPNPNQYKVEFNTPDDQITTPYIV
VVNGHIDPNSKGDLALRSTLYGYNSNIIWRSMSWDNEVAFNNGSGSGDGIDKPVVPEQPDEPGEI
EPIPEDS**A**WSHPQFEK

**Bold**: Amino acids introduced by sub-cloning or remaining after TEV protease-mediated removal of 6xHis tag. Underlined: Strep-tag II

*ClfA-002 (ST5) N2-N3 sequence (after TEV protease removal of N-terminal 6xHis tag)*

**GS**VAADAPAAGTDITNQLTDVKVTIDSGTTVYPHQAGYVKLNYGFSVPNSAVKGDTFKITVPKELNL
NGVTSTAKVPPIMAGDQVLANGVIDSDGNVIYTFTDYVDNKENVTANITMPAYIDPENVTKTGNVTLT
TGIGTNTASKTVLIDYEKYGQFHNLSIKGTIDQIDKTNNTYRQTIYVNPSGDNVVLPALTGNLIPNTKS
NALIDAKNTDIKVYRVDNANDLSESYYVNPSDFEDVTNQVRISFPNANQYKVEFPTDDDQITTPYIV
VVNGHIDPASTGDLALRSTFYGYDSNFIWRSMSWDNEVAFNNGSGSGDGIDKPVVPEQPDEPGEI
EPIPEDS**A**WSHPQFEK

SAR114-Fcs Heavy Chain

QVQLQESGPGLVKPSETLSLTCTVSGGSIQNSYWSWIRQPPGKGLEWIGYLYSSGRTNYTPSLKSR
VTISVDTSKNQFSLKLSSVTAADTAVYYCARTHLGGFHYGGGFWFDPWGQGTLVTVSSASTKGPSV
FPLAPSSKSTSGGTAALGCLVKDYFPEPVTVSWNSGALTSGVHTFPAVLQSSGLYSLSSVVTVPSSSL
GTQTYICNVNHKPSNTKVDKKV*EPKSCDKTHTCPPCP*APEAAGGPSVFLFPPKPKDTLMISRTPEVT
CVVVDVSHEDPEVKFNWYVDGVEVHNAKTKPREEQYQSTYRVVSVLTVLHQDWLNGKEYKCKVS
NKALGAPIEKTISKAKGQPREPQVYTLPPSRDELTKNQVSLTCLVKGFYPSDIAVEWESNGQPENNY
KTTPPVLDSDGSFFLYSKLTVDKSRWQQGNVFSCSVMHEALHNHYTQKSLSLSPGK

Key: CH1, CH2, CH3, positions mutated for Fc silencing: L234A, L235A, N297Q, P329G.
Italics: hinge.

SAR114-Fcs Light Chain

DIQMTQSPSSLSASVGDRVTITCRASQSITSYLNWYQQKPGKAPKLLIYASSSLQSGVPSRFSGSGS
GTDFTLTISSLQPEDFATYYCQESYSTPPTFGQGTKVEIKRTVAAPSVFIFPPSDEQLKSGTASVVCLL
NNFYPREAKVQWKVDNALQSGNSQESVTEQDSKDSTYSLSSTLTLSKADYEKHKVYACEVTHQGL
SSPVTKSFNRGEC

Key: Variable light (VL) domain, human CL kappa

**Figure S1. Protein sequences of ClfA-001, ClfA-002 and SAR114-Fcs.**

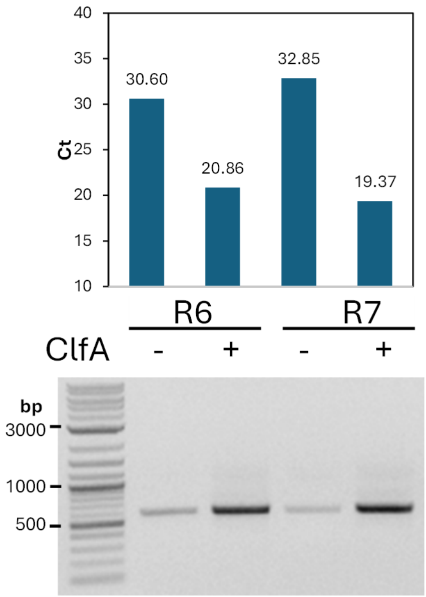


**Figure S2. Pull-down of click-displayed R6/R7 DARPin pools.** Click-displayed DARPin-cDNA was diluted 10-fold in SBTD0.3 buffer and incubated in the absence or presence of 200 nM bClfA at room temperature for 1 hour. Streptavidin-coated magnetic beads were then added to the mixture and the amount of cDNA associated with the beads was quantified using qPCR using the Forget-Me-Not qPCR kit (Biotium) and primers 1892 + 2488. Agarose gel electrophoresis of PCR products is shown beneath the corresponding bar graph.

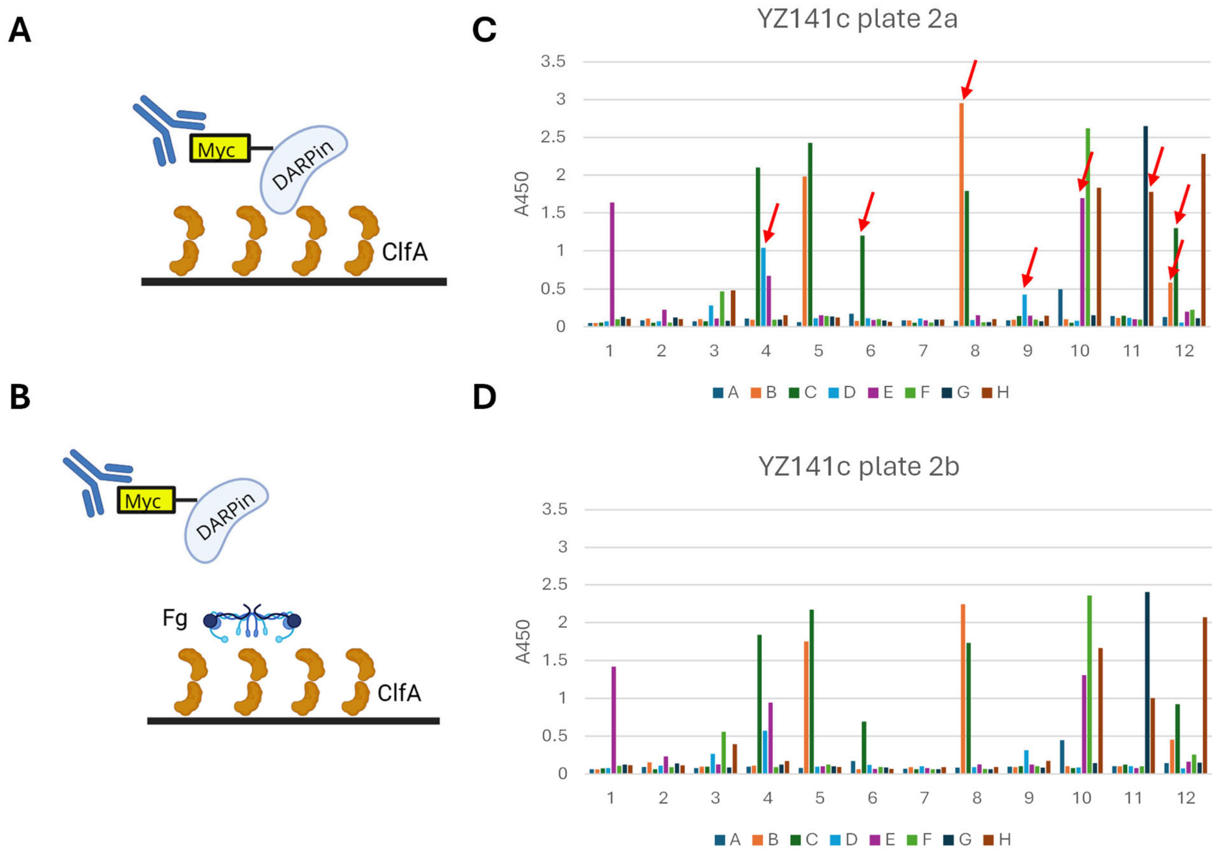


**Figure S3. Functional screens to identify DARPins able to inhibit the Fg-ClfA interaction.** Schematic for the ClfA-binding **(A)** and competition **(B)** ELISA. Representative data from ClfA-binding **(C)** and competition **(D)** ELISA. Red arrows indicate the candidate DARPins selected for further analysis.

| | NCAP / AR1 / AR2 / AR3 / CCAP / |
|---|---|
| | MGSSHHHHHHSSGLVPRGSHMEQKLISEEDLGSDLGKKLLEAARAGQDDEVRILMANGADVNA/XXXXGXTPLHLAAXXGHLEIVEVLLKXGADVNA/XXXXGXTPLHLAAXXGHLEIVEVLLKXGADVNA/XXXXGXTPLHLAAXXGHLEIVEVLLKXGADVNA/XXXXGXTAFDISIDNGNEDLAEILQ |
| D6 | MGSSHHHHHHSSGLVPRGSHMEQKLISEEDLGSDLGKKLLEAARAGQDGEVRILMANGADVNA/SSMHGITPLHLAAARGHLEIVEVLLKHGADVNA/WFRSGPTPLHLAALCGHLEIVEALLKHGADVNA/REWFGFTPLHLAATRGHLEIVEVLLKNGADVNA/QRNTGLTAFDISIDNGNEDLAEILQ |
| D8 | MGSSHHHHHHSSGLVPRGSHMEQKLISEEDLGSDLGKKLLEAARAGQDDEVRILMANGADVNA/RTLDGFTPLHLAAFWGHLEIVEVLLKNGADVNA/TKIYGLTPLHLAAFRGHLEIVEVLLKNGADVNA/RIHPGSTPLHLAASKGHLEIVEVLLKNGADVNA/QLFNGLTAFDISIDNGNEDLAEILQ |
| D13 | MGSSHHHHHHSSGLVPRGSHMEQKLISEEDLGSDLGRKLLEAARAGQDDEVRVLMANGADVNA/YTVLGITPLHLAAFNGHLEIVEALLKDGADVNA/LSAGGVTPLHLAALCGHLEIVEVLLKHGADVNA/REWFGFTPLHLAATRGHLEIVEVLLKNGADVNA/QPPDGNTAFDISIDNGNEDLAEILQ |
| D17 | MGSSHHHHHHSSGLVPRGSHMEQKLISEEDLGSDLGKKLLEAARAGQDDEVRILMANGADVNA/YTVLGITPLHLAAFNGHLEIVEVLLKDGADVNA/LSAGGVTPLHLAALCGHLETVEVLLKHGADVNA/REWFGFTPLHLAATRGHLEIVEVLLKHGAGVNT/QLSNGNTAFDISIDNGDEDLAEILQ |
| D21 | MGSSHHHHHHSSGLVPRGSHMEQKLISEEDLGSDLGKKLLEAARAGQDDEVRILMANGADVNA/ILRFGGTPLHLAAWKGHLEIVEVLLKDGADVNA/GLKYGETPLHLAAVNGHLEIVEVLLKNGADVNA/RKPPGPTPLHLAALTGHSKIVEVLLKNGADVNA/QRNTGLTAFDISIDNGNEDLAEILQ |

X Fully randomized positions

6xHis tag

Myc tag

| | |
|---|---|
| Dx-γ | **Dx-KAAGSPTPTPTTPTPTPTGSGEGQQHHLGGAKQAGDV** |

γ-peptide

PT linker

**Figure S4. Sequence alignment of lead 1st-generation DARPins following eight rounds of click-display selection.**

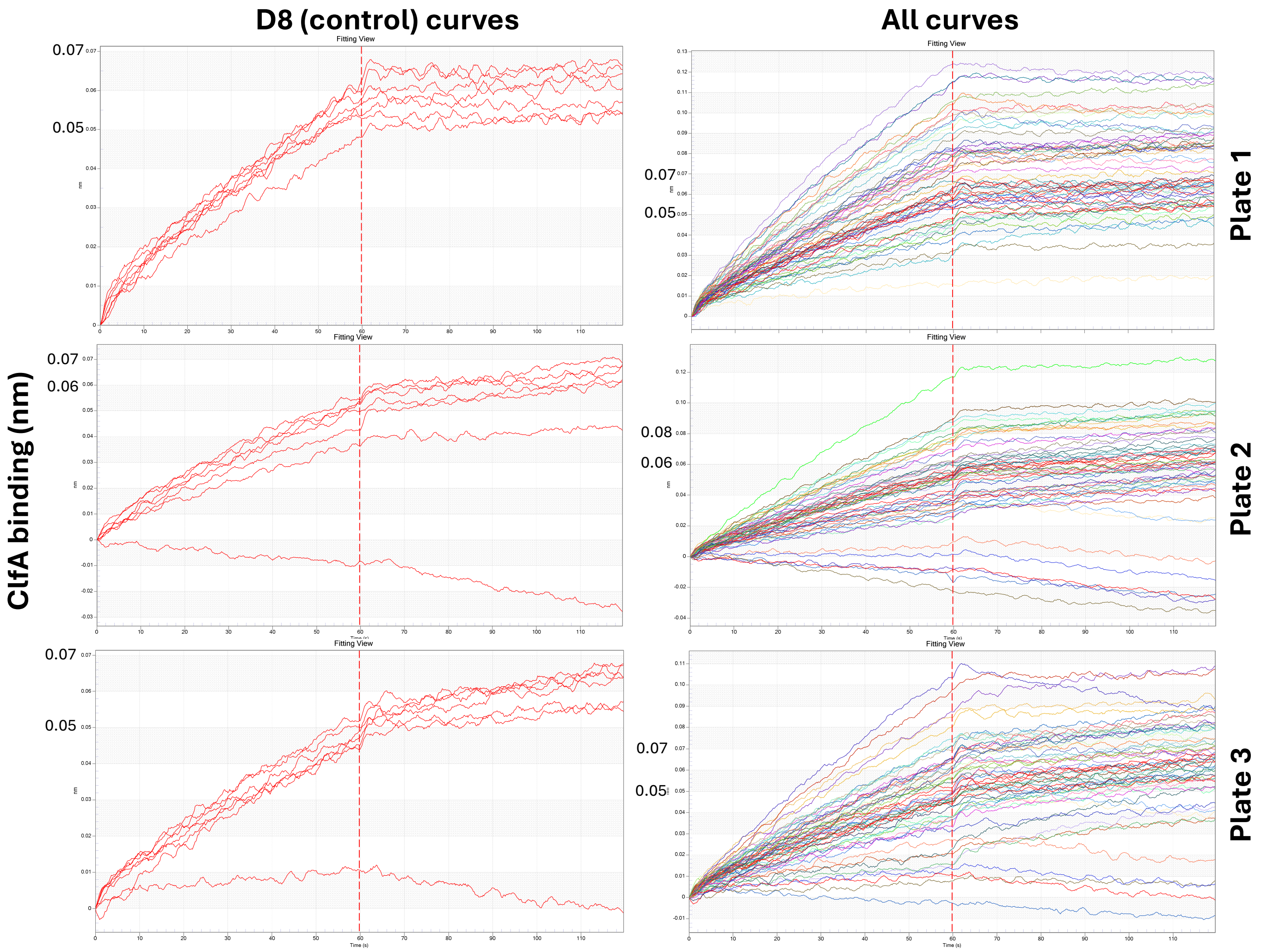


**Figure S5. Affinity maturation of 1st-generation anti-ClfA DARPin D8.** Step-corrected ClfA-binding curves of variants of DARPin D8, present in crude BL21 *E. coli* lysate. The DARPins were sourced from the output of 5 rounds of click-display affinity maturation of D8 following error-prone PCR randomization. Individual BL21 *E. coli* colonies containing DARPins were grown in 1-mL LB medium overnight in deep (2-mL) V-bottom 96-well plates and 10-mL of overnight culture was used to inoculate 1-mL fresh LB prior to growth to mid-log phase ($OD_{600}$ 0.4 – 0.8) and induction with 0.5 mM IPTG at 37°C for 4 h. Harvested cell pellets were resuspended with 200-µL lysis buffer (PBS, 0.2 mg/mL lysozyme, 1 mM $MgCl_2$) and subjected to 3 freeze-thaw cycles (30 mins at -80°C, 20 mins at 37°C). Clarified lysates were diluted 1000-fold with Assay Buffer (PBS, 0.02% Tween-20, 0.1% bovine serum albumin) for BLI analysis. For BLI, a single set of 8 streptavidin (SA) sensors were loaded once with 100 nM biotinylated ClfA (bClfA) in Assay Buffer for 2 mins and re-used for processing all lysate samples as described below. Three 96-well assay plates (Greiner Bio-One # 655209) were each set up with 7 columns containing lysates from 49 variant DARPins (7 candidate affinity-matured D8 lysates per column). One well in each column contained D8 (control) lysate. For each measurement, the 8 bClfA-loaded sensors were first equilibrated in Assay Buffer for 1 min, followed by 1 min association with the diluted lysates, followed by 1 min dissociation in Assay Buffer. The sensors were regenerated via three cycles of the following steps before moving on to measurement of the next set of 8 samples: (i) 5 secs exposure to regeneration buffer (10 mM Glycine-HCl, pH 2.0, 1000 rpm), then (ii) 5 secs exposure to Assay Buffer (1000 rpm). Left, D8 (control) binding curves only; right, all binding curves from the three 96-well screening plates. The y-axis on the different plots is scaled differently, and the binding values bordering the plateau region of the control curves are indicated in larger text to facilitate comparison. The red curves on the right plots designate the D8 control curves. Several clones from each plate showed a higher plateau ClfA-binding response to the parent DARPin D8. Data were generated using an Octet Red96e instrument (ForteBio).

| | $k_D$ (nM) | $k_{on}$ (1/Ms) | $k_{off}$ (1/s) |
|---|---|---|---|
| **ACE6** | 12.06 | 9.825e4 | 1.185e-3 |
| **ACE25** | 12.74 | 9.421e4 | 1.200e-3 |
| **ACE1** | <0.1[†] | 2.905e4 | <1e-7 [†] |
| **ACE3** | <0.1[†] | 3.550e4 | <1e-7[†] |
| **ACE7** | 3.472 | 2.168e4 | 7.528e-5 |
| **ACE10** | <0.1[†] | 1.820e4 | <1e-7[†] |
| **ACE12** | <0.1[†] | 3.110e4 | <1e-7[†] |
| **ACE14** | <0.1[†] | 2.304e4 | <1e-7[†] |
| **D8** | 150.7[††] | 1.13e4[††] | 1.698e-3[††] |

**Figure S6. Binding kinetics of purified lead affinity-matured variants of anti-ClfA DARPin D8 to ClfA.** Fresh BLI sensors were hydrated in Assay Buffer for 10 mins, followed by loading with 100 nM bClfA for 2 mins, then association with 100 nM DARPin variants for 2 mins, then dissociation in Assay Buffer for 2 mins. The selected DARPins were categorized based on their ClfA-binding activity relative to D8 as having either similar off-rate (two clones, top) or low off-rate (six clones, bottom). All eight DARPins showed a higher on-rate and >10-fold lower equilibrium dissociation constant ($k_D$) relative to D8. Shown on the right are the kinetics rates constants computed by the BLItz Pro 1.3 software based on either a local fit of a single association/dissociation curve (candidate affinity-matured D8 variants) or a global fit of three independently generated curves (parent D8 control). Data were generated using a BLItz instrument (ForteBio) and are representative of at least two independent experiments.

[†] These values were lower than the limit of detection of the ForteBio BLItz instrument.

[††] Kinetics constants for the D8 parent (control) were derived from a global fit of 3 independent binding curves.

### (A) Similar off-rate/higher on-rate compared to D8

```
                    N-Cap                             AR1
              1   5    10   15   20   25   30   35   40   45   50   55   60
D8   (parent) GKKLLEAARAGQDDEVRILMANGADVNARTLDGFTPLHLAAFWGHLEIVEVLLKNGADVN
ACE6          GKKLLEAARAGQDDEVRILMANGADVNARTLDGYTPLHLAAFWGHLEIVEVLLKNGADVN
ACE25         GKKLLEAARAGQDDEVRILMANGADVNARTLDGYTPLHLAAFWGHLEIVEVLLKNGADVN
              *********************************:**************************

                    AR2                               AR3
              61   65   70   75   80   85   90   95   100  105  110  115  120
D8 (parent)   ATKIYGLTPLHLAAFRGHLEIVEVLLKNGADVNARIHPGSTPLHLAASKGHLEIVEVLLK
ACE6          ATKIYGLMPLHLAAFRGHLEIVEVLLKNGADVNARIHPGRTPLHLAASKGHLEIVEVLLK
ACE25         ATKIYGLMPLHLAAFRGHLEIVEVLLKNGADVNARIHPGSTPLHLAASKGHLEIVEVLIK
              ******* ******************************* *****************:*

                    C-Cap
              121 125  130  135  140  145  150
D8 (parent)   NGADVNAQLFNGLTAFDISIDNGNEDLAEILQ
ACE6          NGADVNAQLFNGLTAFDISIDNGNEDLAEILQ
ACE25         NGADVNAQLFNGLTAFDIAIDNGNEDLAEILQ
              ******************:*************
```

### (B) Lower off-rate compared to D8

```
              1   5    10   15   20   25   30   35   40   45   50   55   60
D8 (parent)   GKKLLEAARAGQDDEVRILMANGADVNARTLDGFTPLHLAAFWGHLEIVEVLLKNGADVN0
ACE1          GKKLLEAARAGQDGEVRILMANGADVNARTLDGYTPLHLAAFWGHLEIVEVLLKNGADVN60
ACE3          GKKLLEAARAGQDDEVRVLMANGADVNARTLDGYTPLHLAAFWGHLEIVEVLLKNGADVN60
ACE7          GKKLLEAARAGQDDEVRILMANGADVNARTLDGYTPLHLAAFWGHLEIVEVLLKNGADVN60
ACE10         GKKLLEAARAGQDDEVRILMANGADVNARTLDGYTPLHLAAFWGHLEIVEVLLKNGADVN60
ACE12         GKKLLEAARAGQDDEVRILMANGADVNARTLDGYTPLHLAAFWGHLEIVEVLLKNGADVN60
ACE14         GKKLLEAARAGQDDEVRILMANGADVNARTLDGYTPLHLAAFWGHLEIVEVLLKNGADVN60
              *************.***:***************:**************************

              61   65   70   75   80   85   90   95   100  105  110  115  120
D8 (parent)   ATKIYGLTPLHLAAFRGHLEIVEVLLKNGADVNARIHPGSTPLHLAASKGHLEIVEVLLK20
ACE1          ATKIYGLTPLHLAAFRGHLEIVEVLLKNGADVNARIHPGRTPLHLAAWKGHLEVVEVLLK.20
ACE3          ATKIYGLMPLHLAAFRGHLEIVEVLLKNGADVNARIHPGRTPLHLAAWKGHLEVVEVLLK.20
ACE7          ATKIYGLTPLHLAAFRGHLEIVEVLLKNGADVNARIHPGRTPLHLAAWKGHLEVVEVLLK.20
ACE10         ATKIYGLKPLHLAAFRGHLEIVEVLLKNGADVNARIHPGRTPLHLAAWKGHLEIVEELLK.20
ACE12         ATKIYGLKPLHLAAFRGHLEIVEVLLKNGADVNARIHPGRTPLHLAAWKGHLEVVEVLLK.20
ACE14         ATKIYGLKPLHLAAFRGHLEIVEVLLKNGADVNARIHPGRTPLHLAAWKGHLEIVEVLLK.20
              ******* ******************************* ******* *****:** ***

              121 125  130  135  140  145  150
D8 (parent)   NGADVNAQLFNGLTAFDISIDNGNEDLAEILQ   152
ACE1          NGADVNAQLFNGLTAFDISIDNGNEDLAEILQ   152
ACE3          NGADVNAQLFNGLTAFDISIDNGNEDLAEILQ   152
ACE7          NGADVNAQLFNGLTAFDISIDNGNEDLAEILQ   152
ACE10         NGADVNAQLFNGLTAFDISIDNGNEDLAEILQ   152
ACE12         NGADVNAQLFNGLTAFDISIDNGNEDLAEILQ   152
ACE14         NGADVNAQLFNGLTAFDISIDNGNEDLAEILQ   152
              ********************************
```

**Figure S7. Protein sequence alignment of affinity-matured D8-variant DARPins.** Candidates showing a similar off-rate and higher on-rate **(A)** or lower off-rate **(B)** relative to D8 (see Figure 2A of main paper) are separately aligned. The fully randomized positions in the naïve DARPin library are boxed. Mutations relative to the D8 parent are highlighted in cyan (fully conserved within the alignment), yellow (partially conserved, either within a given alignment or across alignments in (A) and (B)), or pink (non-conserved). Shading denotes the AR1, AR2, and AR3 domains (colored) or N-/C-caps (grey).

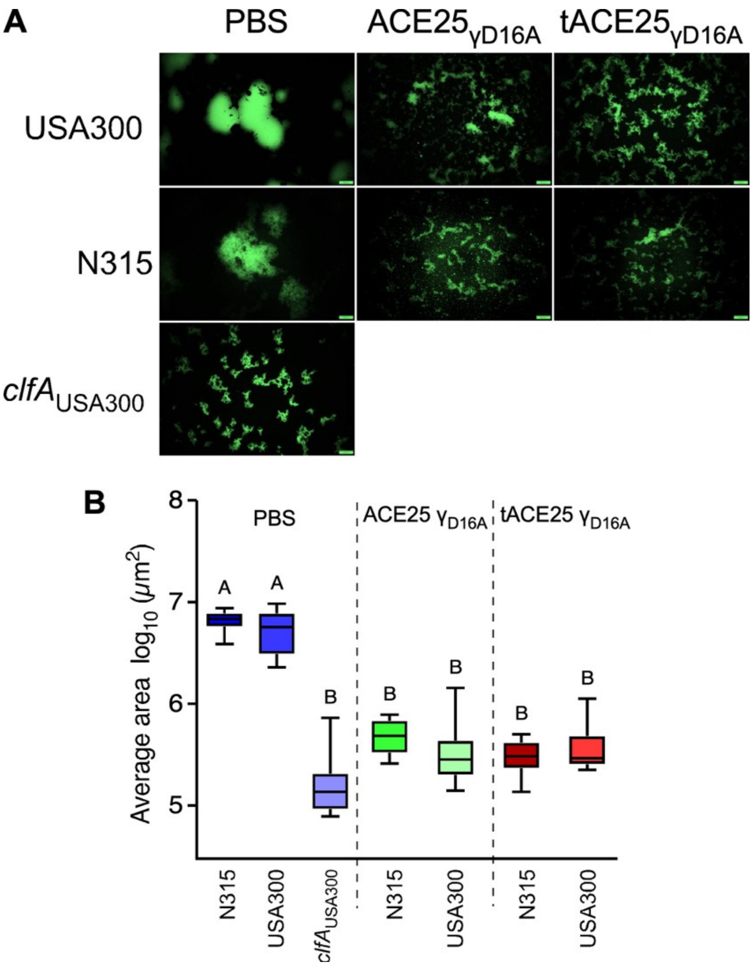


**Figure S8. Staphylococcal agglutination in human plasma.** (A) Representative images of SYTO-9-stained two serotypes of *S. aureus* (USA300-LAC and N315) bacteria agglutinated in human plasma with anti-ClfA DARPins or PBS (negative treatment). *clfA* isogenic mutant was used as positive control. Images were observed under an inverted fluorescence microscope with a 20× objective. (B) Box and whisker plot representation of staphylococcal agglutination areas with anti-coagulated human plasma observed in 10 different fields of microscopic view. Statistical significance was calculated in pairwise comparison using Brown-Forsythe and Welch ANOVA followed by Dunnett's T3 correction for multiple comparisons. Letters denote statistical significance as observed by multiple comparisons across samples. Experiment was performed twice for reproducibility.

CADAR1 protein sequence (1 copy of serum-binding DARPin @ N terminus)

MGHHHHHHGEQKLISEEDLENLYFQGSDLGKKLLEAARAGQDDEVRELLKAGADVNAKDYFSHT
PLHLAARNGHLKIVEVLLKAGADVNAKDFAGKTPLHLAANEGHLEIVEVLLKAGADVNAQDIFGKTP
ADIAADAGHEDIAEVLQKAAGSPTPTPTTPTPTPTTPTPTPTGSKKKPLDGEYFTLQIRGRERFEMFRE
LNEALELKDAQAGKEPGGGGGSGGGGSDLGKKLLEAARAGQDDEVRILMANGADVNARTLDGY
TPLHLAAFWGHLEIVEVLLKNGADVNATKIYGLMPLHLAAFRGHLEIVEVLLKNGADVNARIHPGST
PLHLAASKGHLEIVEVLIKNGADVNAQLFNGLTAFDIAIDNGNEDLAEILQKAAGSPTPTPTTPTPTPT
GSGEGQQHHLGGAKQAGAV

- 6xHis tag
- Myc tag
- TEV Cleavage site
- DA50 (serum-albumin-binding DARPin)
- Long PT linker (20 aa)
- p53-TD
- Flexible linker
- DARPin ACE25
- Short PT linker (13 aa)
- Fg-$\gamma_{D16A}$ peptide

CADAR2 protein sequence (2 copies of serum-binding DARPin @ N terminus)

MGHHHHHHGEQKLISEEDLENLYFQGSDLGKKLLEAARAGQDDEVRELLKAGADVNAKDYFSHT
PLHLAARNGHLKIVEVLLKAGADVNAKDFAGKTPLHLAANEGHLEIVEVLLKAGADVNAQDIFGKTP
ADIAADAGHEDIAEVLQKAAGSPTPTPTTPTPTPTTPTPTPTGSDLGKKLLEAARAGQDDEVRELLKA
GADVNAKDYFSHTPLHLAARNGHLKIVEVLLKAGADVNAKDFAGKTPLHLAANEGHLEIVEVLLKA
GADVNAQDIFGKTPADIAADAGHEDIAEVLQKAAGSPTPTPTTPTPTPTTPTPTPTGSDLGKKLLEAAR
AGQDDEVRILMANGADVNARTLDGYTPLHLAAFWGHLEIVEVLLKNGADVNATKIYGLMPLHLAAF
RGHLEIVEVLLKNGADVNARIHPGSTPLHLAASKGHLEIVEVLIKNGADVNAQLFNGLTAFDIAIDNG
NEDLAEILQKAAGSPTPTPTTPTPTPTGSGEGQQHHLGGAKQAGAV

- 6xHis tag
- Myc tag
- TEV Cleavage site
- DA50 (serum-albumin-binding DARPin)
- Long PT linker (20 aa)
- Flexible linker
- DARPin ACE25
- Short PT linker (13 aa)
- Fg-$\gamma_{D16A}$ peptide

**Figure S9. Protein sequences of 2$^{nd}$-generation DARPin constructs, including a serum-albumin-binding DARPin (DA50) component.**

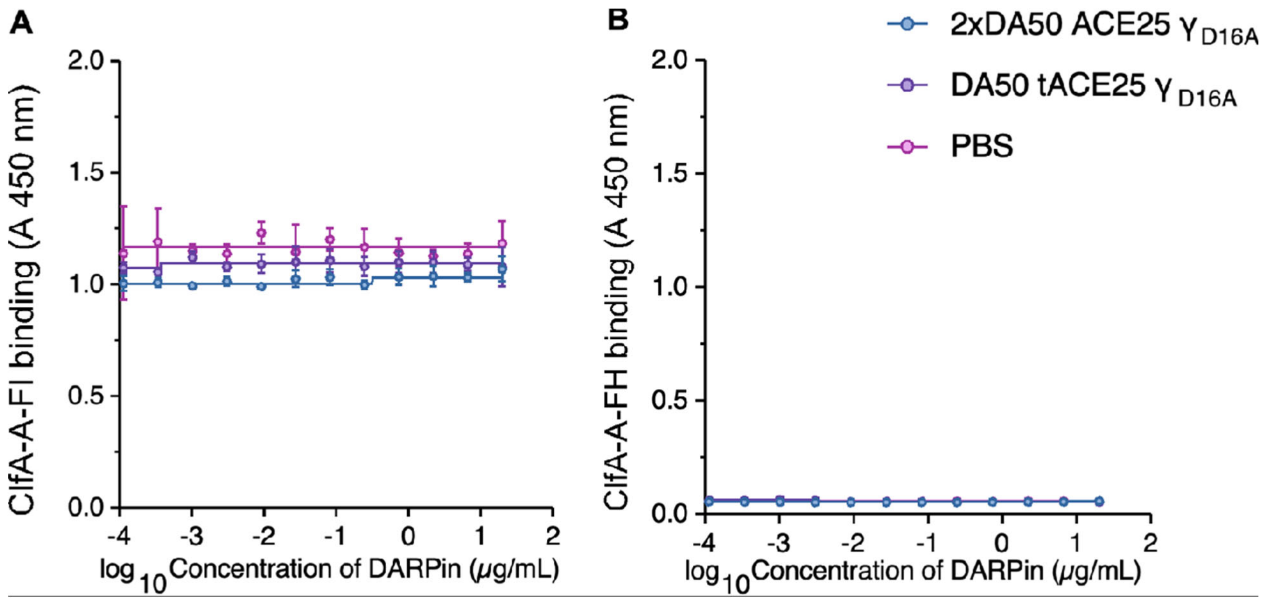


**Figure S10. Competitive ELISA experiment between human fI, fH and DARPins with ClfA-A.** Human fI **(A)** or fH **(B)** were added to ELISA plates coated with ClfA-A from Newman. Serial dilution of DARPins were added as a competitor to the wells. PBS served as background control. Bound factor I or factor H were detected using anti fI or anti fH primary antibody followed by goat anti-rabbit HRP labeled secondary antibody. Absorbance at 450 nm was recorded to determine binding. Experiments were performed twice for reproducibility.

*Conventional selection*

| Round | *In vitro* translation scale† | Negative selection on empty Streptavidin beads? | bClfA target (nM) | Library-bClfA incubation time (min) | Washes |
|---|---|---|---|---|---|
| 1 | Full | No | 20 | 60 | 6 |
| 2 | Half | Yes | 10 | 60 | 8 |
| 3 | Half | Yes | 2.5 | 10 | 12†† |

*Off-rate selection*

| Round | *In vitro* translatio n scale† | Negative selection on empty Streptavidin beads? | bClfA target (nM) | Library-bClfA incubation time (min) | Fg incubation time | Washes |
|---|---|---|---|---|---|---|
| 4 | Half | Yes | 1 | 5 | 1 h (500 nM Fg) at RT | 12†† |
| 5 | Half | Yes | 1 | 1 | (i) Overnight at RT with 500 nM Fg, then (ii) 4 h at RT and 2 h at 37°C with 1000 nM Fg | 12†† |

†Full-scale *in vitro* translation reactions (PURExpress *In Vitro* Protein Synthesis Kit, NEB) for the generation of click-displayed protein libraries utilized 1 µg template DNA and a 25-µL final reaction volume. Half-scale utilized 0.5 µg template DNA and 12.5-µL final reaction volume.
††The beads were incubated with biotinylated filler protein for 30 minutes at room temperature prior to the 7th wash.
bClfA: biotinylated ClfA(ST8)

**Table S1. Summary of parameters employed during affinity maturation of anti-ClfA DARPin D8 by click display.** Three rounds of conventional click-display selection were followed by two rounds of off-rate selection.

### List of primers used in this study

**Primers used for amplification of click-display DARPin template**
1938: ATA CGA AAT TAA TAC GAC TCA CTA GGA GAC CAC AAC GGT TTC CCT CTA GAA ATA ATT TTG TTT AAC
2643: AAA CCC CTC CGT TTA GAG AGG GGT TAT GCT AGT TAT TCC ACG CCG CCC CCC GTC CT

**Primers used for qPCR detection of click-displayed DARPins**
1892: TTG AGG AGA GTA GGA CCT
2488: GTT TAA CTT TAA GAA GGA GGA TAT ATC CAT
